# Influence of Arm Swing on Cost of Transport during Walking

**DOI:** 10.1101/426775

**Authors:** Myriam L. de Graaf, Juul Hubert, Han Houdijk, Sjoerd M. Bruijn

## Abstract

Normal arm swing plays a role in decreasing the cost of transport during walking. However, whether excessive arm swing can reduce the cost of transport even further is unknown. Therefore, we tested the effects of normal and exaggerated arm swing on the cost of transport in the current study. Healthy participants (*n*=12) walked on a treadmill (1.25 m/s) in seven trials with different arm swing amplitudes (in-phase, passive restricted, active restricted, normal, three gradations of extra arm swing), while metabolic energy cost and the vertical angular momentum (VAM) and ground reaction moment (GRM) were measured.

In general, VAM and GRM decreased as arm swing amplitude was increased, except for in the largest arm swing amplitude condition. The decreases in VAM and GRM were accompanied by a decrease in cost of transport from in-phase walking (negative amplitude) up to a slightly increased arm swing (non-significant difference compared to normal arm swing). The most excessive arm swings led to an increase in the cost of transport, most likely due to the cost of swinging the arms. In conclusion, increasing arm swing amplitude leads to a reduction in vertical angular moment and ground reaction moments, but it does not lead to a reduction in cost of transport for the most excessive arm swing amplitudes. Normal or slightly increased arm swing amplitude appears to be optimal in terms of cost of transport in young and healthy individuals.

**SUMMARY STATEMENT:** Excessive arm swing reduces the vertical angular momentum and ground reaction moment, but not necessarily the energetic cost of transport.

## INTRODUCTION

Human locomotion distinguishes itself from that of many other vertebrates due to its predominantly two-legged nature. Therefore, it is not surprising that most research into human locomotion focuses on the lower extremities, while the contribution of the upper extremities is neglected. However, the arms do appear to play a significant role in locomotion. Studies have shown that people consume more energy when they do not swing their arms during walking (Collins et al., 2009a; Umberger, 2008). This indicates a cost-reducing function of arm swing. There is also evidence that arm swing may be involved in regulating the stability of locomotion (Meyns et al., 2013).

How arm swing is instigated during walking is not yet fully known. Some studies mention a predominantly passive nature, as a result of the dynamics of the linked body segments (Collins et al., 2009a; Gerdy, 1829; Jackson et al., 1978; Morton and Fuller, 1952; Weber and Weber, 1836), where the arms would then function as passive pendulums. Other studies brought this idea into question as they found muscle activity in the upper extremities (Fernandez-Ballesteros et al., 1965; Kuhtz-Buschbeck and Jing, 2012), thereby indicating an active origin of arm swing. However, the muscle activity is also present during walking when the arms are bound at the sides (Kuhtz-Buschbeck and Jing, 2012), and contains co-contraction of two agonistic parts of the deltoid (Pontzer et al., 2009) which points at either activation through central pattern generators (see Zehr & Duysens (2004)) or a more stabilizing function (Meyns et al., 2013). It has also been found that passive dynamics are sufficient to generate arm swing (Collins et al., 2009a; Jackson et al., 1978), but that the resulting amplitude and relative phase decrease significantly without muscle activity (Goudriaan et al., 2014). Together these findings seem to indicate a role for both active and passive components in the generation of arm swing amplitude.

Independent of how arm swing is executed, it appears to play an important part during human locomotion. However, what this role is exactly, is still unknown. Several hypotheses have been formulated, among which: (a) reducing vertical displacement of the center of mass (COM) (Hinrichs, 1990; Murray et al., 1967; Pontzer et al., 2009; Umberger, 2008), (b) reducing angular momentum around the longitudinal axis (Bruijn et al., 2008; Bruijn et al., 2011; Collins et al., 2009a; Elftman, 1939; Hinrichs, 1990; Park, 2008); (c) reducing angular movement around the longitudinal axis (Fernandez-Ballesteros et al., 1965; Murray et al., 1967; Pontzer et al., 2009); (d) reducing the ground reaction moment (GRM) (Collins et al., 2009a; Li et al., 2001; Witte et al., 1991); (e) increasing (local) stability (Ortega et al., 2008) / balance recovery after perturbations (Bruijn et al., 2010; Hof, 2007; Marigold et al., 2002; Pijnappels et al., 2010); (f) facilitating leg movement (Meyns et al., 2013) and; (g) minimizing energetic costs (Collins et al., 2009a; Ortega et al., 2008; Umberger, 2008). These hypotheses cannot be seen entirely separate from each other, and are in some cases even entirely interdependent.

This study focusses on the relevance and interplay of three of the roles mentioned above, namely those in energetic cost, vertical angular momentum (VAM) and ground reaction moments (GRM). Arm swing is often viewed as a mechanism to decrease angular momentum of the whole body around the vertical axis (as mentioned in hypothesis b above). This idea is based on the observation that angular momentum of the arms is fairly equal in size, but opposite in direction to the momentum of the body (Elftman, 1939; Herr and Popovic, 2008). The change in VAM that results from leg action during walking can, therefore, be compensated by an opposite change in angular momentum through arm swing, thereby bringing the VAM closer to zero (Hinrichs, 1990). The direction of the VAM changes sign during double support, in preparation for the next step. This redirection can be carried out through the legs via the GRM, or, it can be (partially) performed through arm swing: when whole-body VAM is decreased through arm swing, the GRM that needs to be generated by the legs to redirect the VAM will be smaller. With that, the GRF and the forces that the legs need to generate will also be smaller (that is, if stride length and step width – both determining factors for the GRF moment arms – remain unchanged). By this action, a reduction of the whole-body VAM via the arm swing can lead to a decreased energy expenditure by the legs, because the leg muscles do not need to generate as large a GRM. If this were to lead to a decreased total energy expenditure, the energy gain at the legs should be greater than a potential increase in energy expenditure by the arms, i.e. the arms should be more efficient in redirecting the VAM than the legs. This could indeed be the case, because of the suspected (largely) passive nature of arm swing that was discussed before.

Multiple studies have shown that normal arm swing indeed leads to a reduced VAM, when compared to walking without arm swing (Bruijn et al., 2008; Collins et al., 2009a; Elftman, 1939; Herr and Popovic, 2008; Hinrichs, 1990; Park, 2008), with an accompanying reduction in energy expenditure (Collins et al., 2009a; Ortega et al., 2008; Umberger, 2008). Since the VAM is not equal to zero during normal walking, an increase in arm swing amplitude could further decrease VAM. Whether this would then lead to a further reduced cost of transport is unknown. The arm muscles will likely need more activation to increase arm swing amplitude, where normal arm swing appears to be largely passive in nature (Gerdy, 1829; Kubo et al., 2004; Pontzer et al., 2009).

There are studies that have investigated the effect of arm swing amplitude on VAM and energetic costs during walking (e.g. Collins et al. (2009a)). To our knowledge, none of these have looked at arm swing with an amplitude larger than in normal walking. Including extra arm swing conditions can provide extra insight into the (mechanisms behind the) potential energetic cost reducing function of arm swing. Such insight could prove beneficial in multiple situations, e.g. people attempting to lose weight might prefer to use more energy while walking, while elite racewalkers or patients with an increased cost of transport might benefit from energy reducing adaptations.

This study aimed to clarify the relationship between arm swing amplitude and the energetic cost of walking, as well as the role of VAM and GRM herein. We hypothesized that: (1) when arm swing amplitude increases, VAM decreases, (2) when arm swing amplitude increases, GRM decreases, (3) a lower absolute VAM is accompanied by a lower energetic cost, and (4) a lower absolute GRM is accompanied by a lower energetic cost. We defined arm swing amplitude as the difference between the anteroposterior COM position of the two arms, where an arm swing in anti-phase with the legs leads to a positive amplitude and an arm swing in-phase with the legs leads to a negative amplitude. With the resulting information, we hope to give a comprehensive answer on the influence of arm swing on the energetic cost of transport during walking.

## METHODS

### Participants

Twelve healthy subjects have been included in this study (see Table 1). The number of participants included was based on previous studies with a similar design (e.g. (Collins et al., 2009b)). Exclusion criteria were: any orthopedic or neurological disorders that impede gait, and an inability to walk for 5 straight minutes. The experiment was approved by the local ethical committee (Scientific and Ethical Review Board (VCWE), protocol VCWE-2017-040). Prior to the trials, all the participants were informed about the measurements and all signed the informed consent. Participants were free to ask questions at any time and to stop the test if needed.

**Table 1.**
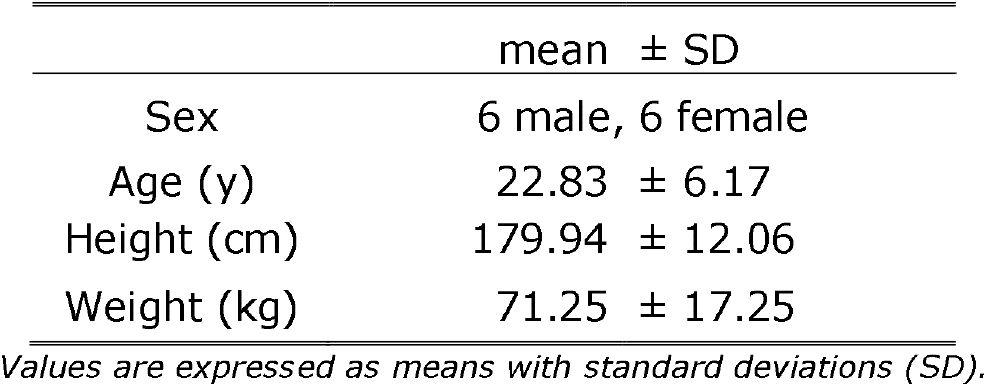
Participant characteristics.

| | mean ± SD |
| --- | --- |
| Sex | 6 male, 6 female |
| Age (y) | 22.83 ± 6.17 |
| Height (cm) | 179.94 ± 12.06 |
| Weight (kg) | 71.25 ± 17.25 |

*Values are expressed as means with standard deviations (SD).*

### Experiment

All participants executed the seven trials, each lasting five minutes, while walking at a speed of 1.25 m/s on a Dual Belt Treadmill. They used a different arm swing amplitude for each trial: (1) normal, (2) held, (3) bound, (4) in-phase, (5) extra I, (6) extra II, (7) extra III. Trials were performed in randomized order, with a few exceptions: the normal condition was always done first to prevent conscious thoughts about arm swing from influencing the normal walking pattern, and the three extra conditions were always performed consecutively from smallest (*extra I*) to largest (*extra III*). Prior to starting the measurements, the participants performed a practice trial to get used to the treadmill and equipment. Verbal instructions and a demonstration of the arm swing that had to be performed were given before each trial: (1) in the *normal* condition participants were told to walk like they always do; (2) in the *held* condition, participants held their arms straight along their body to prevent them from swinging; (3) the *bound* condition was similar to condition 2, only now the arms were bound to the waist with Velcro straps; (4) in the *in-phase* condition the participants were instructed to move their left arm forward with the left leg, and the right arm with the right leg; (5) in *extra I*, the participants were told to increase arm swing slightly as compared to normal, about 1/3 between normal arm swing and the horizontal; (6) In *extra II*, the arm swing had to be at about 2/3 between the horizontal and normal arm swing; (7) in the *extra III* condition, participants were instructed to raise their leading arm up to the horizontal, i.e. parallel to the ground. The instructions for the three extra conditions were given at the same time, to allow the participant to compare the three amplitudes. Adherence to the conditions was visually monitored by the researchers and the participants were told to correct the arm swing when necessary. Participants had the opportunity to take a break after each trial.

### Measurements

We measured: (1) respirometry data with a Cosmed Quark B2 (Cosmed BV, Italy) breath-by-breath respirometer, (2) kinematic data with 17 sensor Xsens MVN inertial sensor suit, sampled at 120 Hz (Xsens Technologies BV, Enschede, The Netherlands), and (3) kinetic data with force sensors in the Dual Belt Treadmill (Y-Mill, ForceLink B.V., The Netherlands) at 1000 Hz.

### Data Analysis

First, separate step cycles were identified by determining left heel strikes on the basis of local minima in the vertical position of the left heel. Then, to check whether the participants had followed the instructions, the arm swing amplitude was analyzed. This was done by calculating the Centre of Mass (COM) position 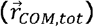 for each arm, using Eqn. 1 with the upper arm, lower arm, and hand segments (s=3).

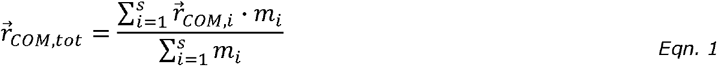

Arm swing amplitude was then calculated as the anteroposterior distance between the COM of the left arm and the COM of the right arm at every point in time. The peak arm swing amplitude of every stride was determined and averaged per condition. The arm swing amplitude was manually made negative for the in-phase condition after we did a visual check to see whether the arm swing really was in-phase with the legs.

#### Cost of Transport

To ensure steady state we only used respirometry data collected during the last two minutes of each trial for the energetic cost calculations. The energy consumption 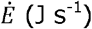 was calculated from the oxygen consumption (VO_2_) and the respiratory exchange ratio (RER) using Eqn. 2 (Garby and Astrup, 1987).

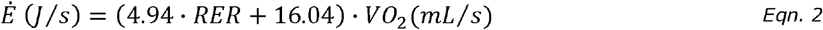

Hereafter, the energy consumption was normalized for body weight and speed to get the cost of transport in J kg^−1^ m^−1^.

#### Vertical Angular Momentum

We calculated total-body COM using equation 1 with all segments. Next, we calculated the VAM around the center of mass as:

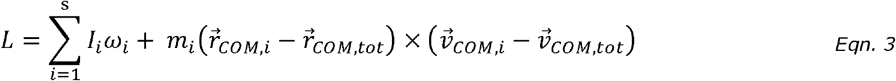

where *L* is the total-angular momentum, *I*_*i*_ is the inertia tensor of segment, *w*_*i*_ is the angular velocity, *m* is the mass, *r* are the position vectors, *v* is velocity. Since the term *I*_*i*_ ⋅ *w*_*i*_ is very small, we have ignored it in the current study. Contributions of the arms and legs to the total, whole-body VAM were also calculated. For this, only the three relevant segments for each extremity were input in the equations. The absolute mean values of the angular momenta per stride were expressed as a measure of the VAM magnitude for all conditions. Using the heel strike indices, VAM was also expressed as a function of the gait cycle, in order to gain a better understanding of the development of direction and magnitude of the VAM during an average stride.

#### Ground Reaction Forces and Moments

Data from the Dual Belt force platform were filtered with a 20 Hz, 2^nd^ order Butterworth filter. The ground reaction moment (GRM) is the moment around the vertical axis caused by the interaction between the feet and floor and knows two components: a pure moment under each individual foot (present during single and double stance), and a pure moment resulting from the force couple created by the horizontal ground reaction forces of both feet (only present during double stance). The ground reaction moment was calculated from the ground reaction force using the following equation (Li et al., 2001):

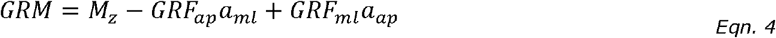

In this formula, *GRM* is the ground reaction moment around the vertical axis, M_z_ is the total vertical moment around the origin of the force platform, *GRF* is the measured ground reaction force, and *a* is the distance between the origin and the COP of the force platform. We calculated the mean absolute GRM value per stride as well as the mean cycle for every condition.

#### Step Parameters

Spatiotemporal parameters can be freely chosen by the participants, meaning they can differ between trials. Therefore the step parameters have been analyzed as indicators to assess how the gait pattern changes as a result of the changes in arm swing amplitude. Step width was calculated as the mean difference between the minimal and maximal x-coordinate of the COP position per gait cycle. Stride frequency was calculated from the time difference between subsequent left heel strikes.

### Statistics

Statistical analysis was performed using IBM SPSS Statistics 23. First, a repeated measures analysis of variance (RM-ANOVA, *α*=0.05) was performed on the arm swing amplitude to test if the conditions indeed lead to the expected behavior and differed between conditions. Then, to test how arm swing (i.e. condition) affects energetics, repeated measures ANOVA was performed on the mean energy costs 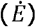. Lastly, to see how arm swing affected kinetics (i.e. GRF and VAM), RM-ANOVAs were performed on the VAM and GRM. To test other possible influences, analysis of variance was also done for step width and step length. If there was a significant main effect, a post hoc paired t-test with Bonferroni correction was executed. Mauchly’s test was used to test for violations of sphericity. If Mauchly’s test was significant, the Greenhouse Geisser corrected values were reported.

### Data Availability

All data, as well as the software used to analyze it, has been made available online. All files can be downloaded from: https://doi.org/10.5281/zenodo.2671651.

## RESULTS

All participants (*n*=12) successfully performed the 7 trials. Oxygen data was compromised in the first three participants, so the cost of transport has only been evaluated in 9 participants. For one participant (#9), oxygen uptake data had to be redone at a later time. The experimental manipulation was successful, as clear effects of condition on arm swing amplitude were found (effect of condition, *F*_*condition*_(2.23, 24.54)=130.04, *p*<.001, see Fig. 1). Post-hoc Bonferroni analysis showed that all conditions differed significantly from each other (*p*≤.001) except for the three extra conditions amongst themselves, *p*>.05).

**Fig. 1.**
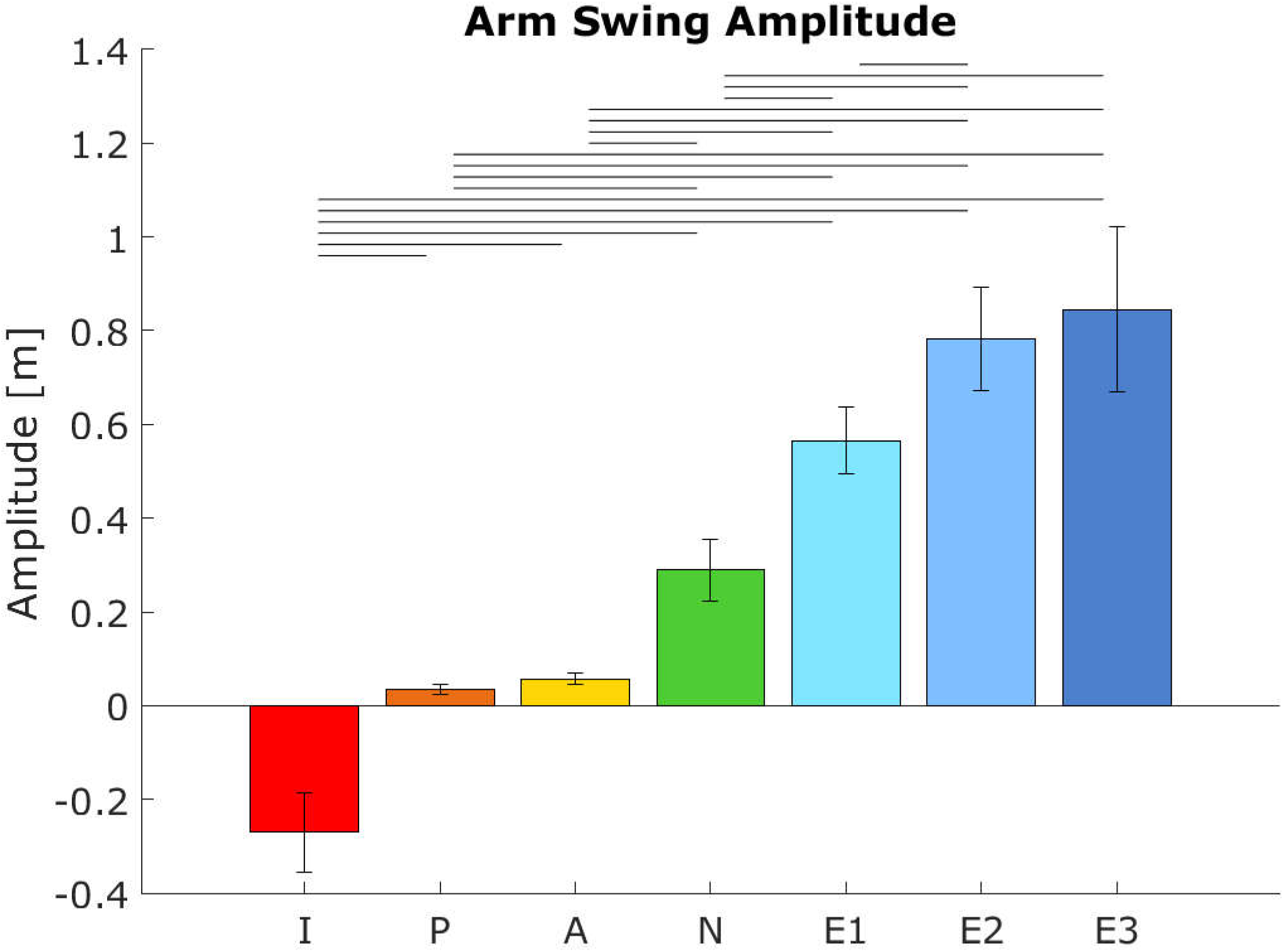
Mean arm swing amplitude (m) per condition. Error bars indicate the 95% Confidence Interval. Horizontal bars show significant differences between conditions (n=12, p<.05, paired t-test with bonferroni and Greenhouse Geisser correction). Abbreviations: I=in-phase, P=passive (restricted), A=active (restricted), N=normal, E=extra.

### Cost of Transport

The cost of transport was higher in conditions with a smaller arm swing amplitude (F_condition_(6,48)=11.95, *p*<.001, see also Fig. 2). Post-hoc analysis showed that in-phase and *extra III* had a significantly higher cost of transport compared to normal (respectively +15.3% and +17.5%, *p*<.05). *In-phase* the cost of transport was also significantly higher than *passive*, *extra I* and *extra II* (*p*<.05), and *extra III* also had a higher cost of transport compared to both other *extra* arm swing conditions.

**Fig. 2.**
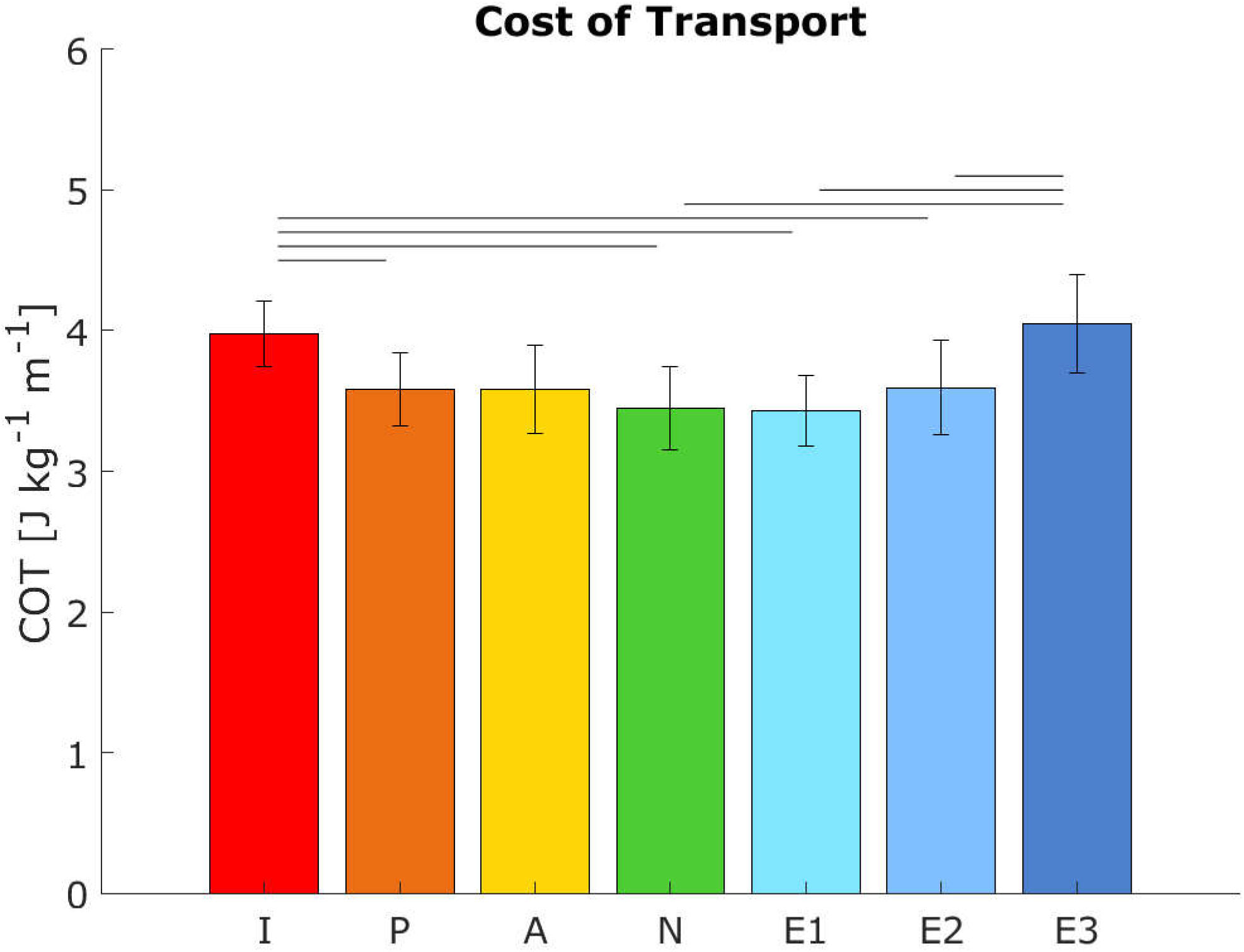
The mean cost of transport (J kg-1 m-1) per condition. Error bars indicate the 95% Confidence Interval. Horizontal bars show significant differences between conditions (n=9, p<.05, paired t-test with bonferroni correction). Abbreviations: I=in-phase, P=passive (restricted), A=active (restricted), N=normal, E=extra.

### Vertical Angular Momentum

The conditions with a lower arm swing amplitude yielded higher whole-body VAM values (*F*_*condition*_(2.67, 29.37)=21.70, *p*<.001, see also Fig. 3A-B). Post-hoc analysis showed significantly higher VAM than normal in *in-phase* (+88.45%, *p*=.008), *passive* (+53.64%, *p*<.001) and *active* (+56.78%, *p*<.001) and significantly lower VAM than normal in *extra II* (−28.16%, *p*=.03). The VAM in the conditions *extra I* and *extra III* were non-significantly lower than normal (respectively −16.61% and −7.98%, *p*>.05).

**Fig. 3.**
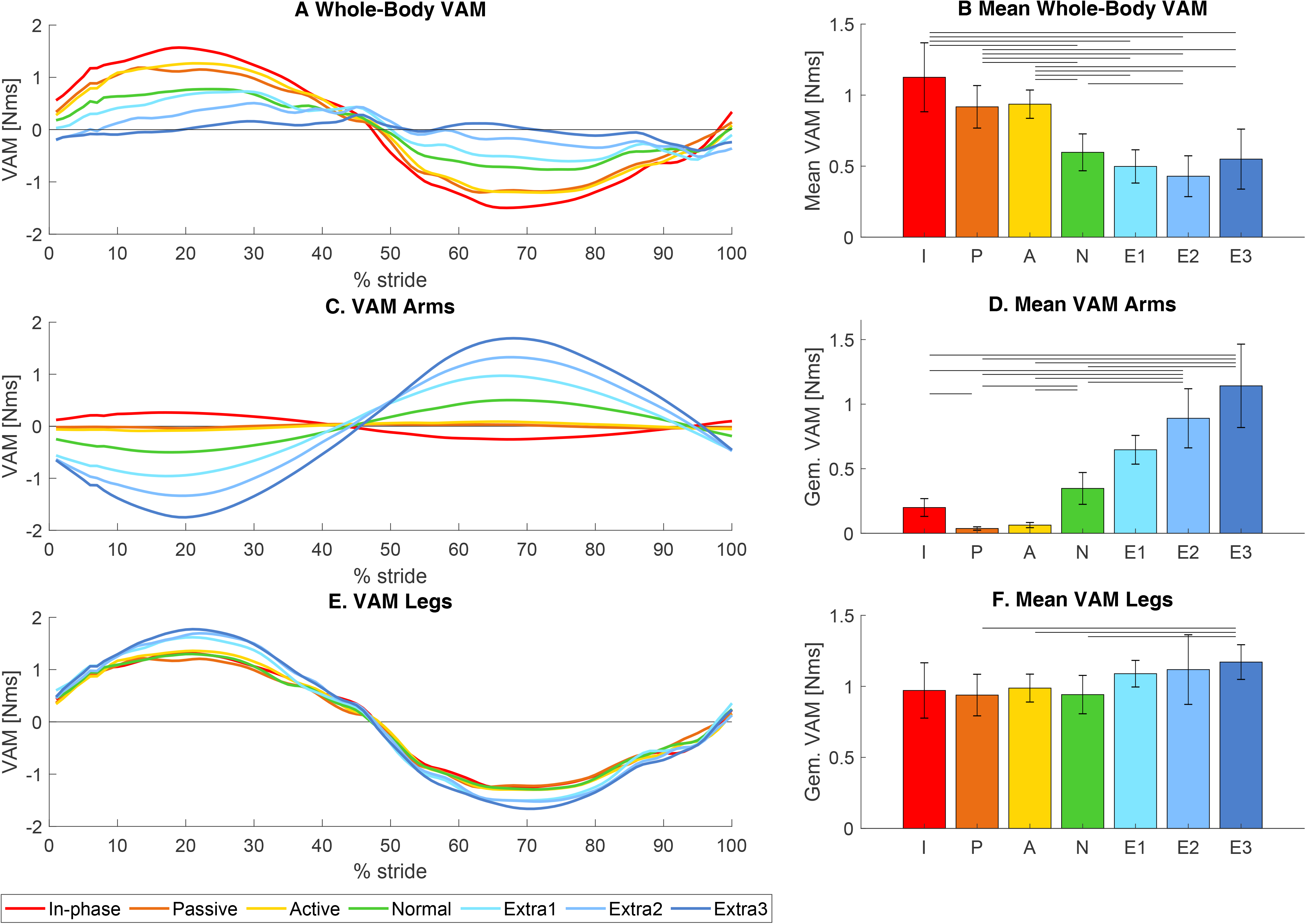
Vertical angular momentum. (A, C, and E) show the mean VAM for all seven trials as a function of the gait cycle (starting and ending with left heel strike), for whole-body VAM, VAM originating from the arms, and VAM originating from the legs respectively. (B, D, and F) show the mean absolute values for whole body-VAM and VAM originating from the arms and legs. Error bars indicate the 95% Confidence Interval. Horizontal bars show significant differences between conditions (n=12, p<.05, paired t-test with bonferroni and Greenhouse Geisser correction). Abbreviations: I=in-phase, P=passive (restricted), A=active (restricted), N=normal, E=extra.

Apart from looking at the whole-body VAM, the contributions of the arms and the legs can be quantified separately as well. The VAM of the arms was significantly higher for conditions with higher arm swing amplitudes (*F*_*condition*_(2.16, 23.80)=38.59, *p*<.001, see also Fig. 3C-D). The VAM of the legs fell just short of a significant relation (*F*_*condition*_(2.27, 25.01)=3.00, *p*=.062, see also Fig. 3E-F).

### Ground Reaction Moments

The conditions with a lower arm swing amplitude had higher GRM values (*F*_*condition*_(1.61, 17.74)=25.69, *p*<.001, see also Fig. 4). Post-hoc analysis showed significantly higher GRM than normal during *in-phase* (+53.62%, *p*=.033) and *passive* (+21.33%, *p*<.001) and *active* (+15.64%, *p*=.004), and significantly lower VAM than normal during *extra II* (−21.96%, *p*=.013) and *extra III* (−25.68%, *p*=.044). The GRM in *extra I* was non-significantly lower than normal (−12.03% *p*>.05).

**Fig. 4.**
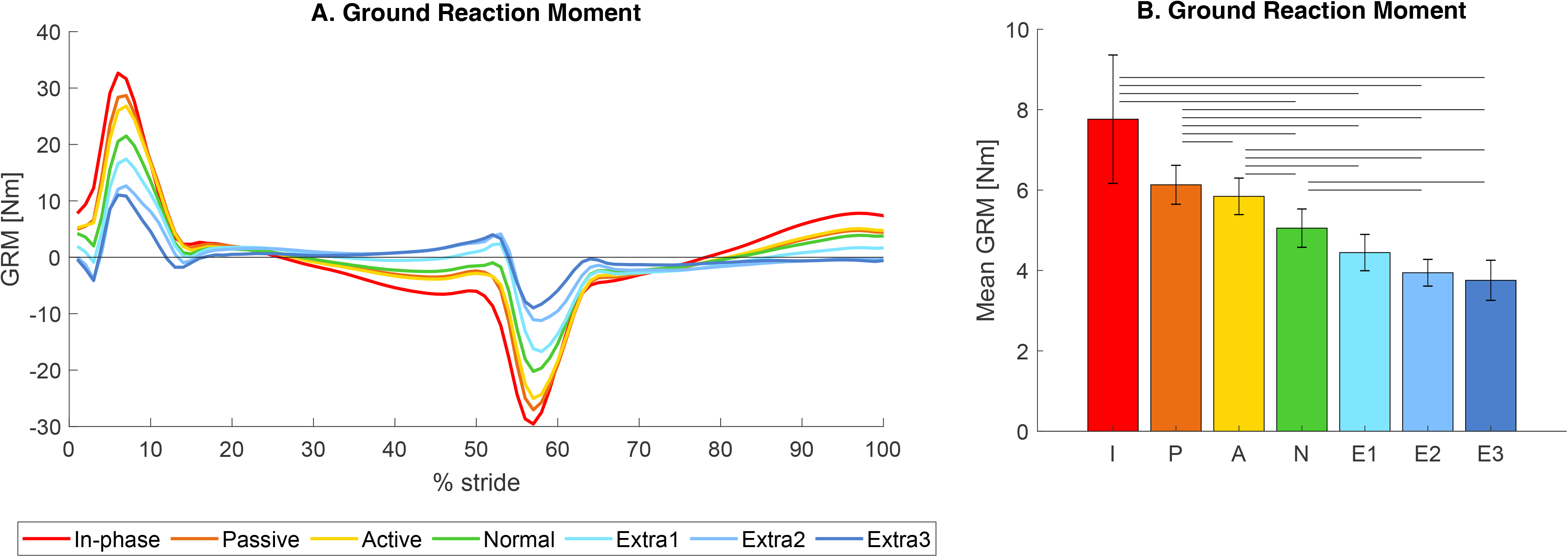
Ground Reaction Moment. (A) shows the mean GRM for all seven trials over one gait cycle from left heel strike to the next heel strike. (B) Shows the mean absolute GRM averaged over all participants per condition. Error bars indicate the 95% Confidence Interval. Horizontal bars show significant differences between conditions (n=12, p<.05). Abbreviations: I=in-phase, P=passive (restricted), A=active (restricted), N=normal, E=extra.

### Step Parameters

Several step parameters have also been analyzed (see Fig. 5). Step width differences over the conditions showed a similar pattern as the ML GRF. There was a significant effect of condition for the step width (*F*_*condition*_(1.92, 21.14)= 4.91, *p*=.005), with post-hoc differences between *active* and the three extra arm swing conditions (all *p*<.05) as well as between *passive* and *extra III* (*p*=.002).

**Fig. 5.**
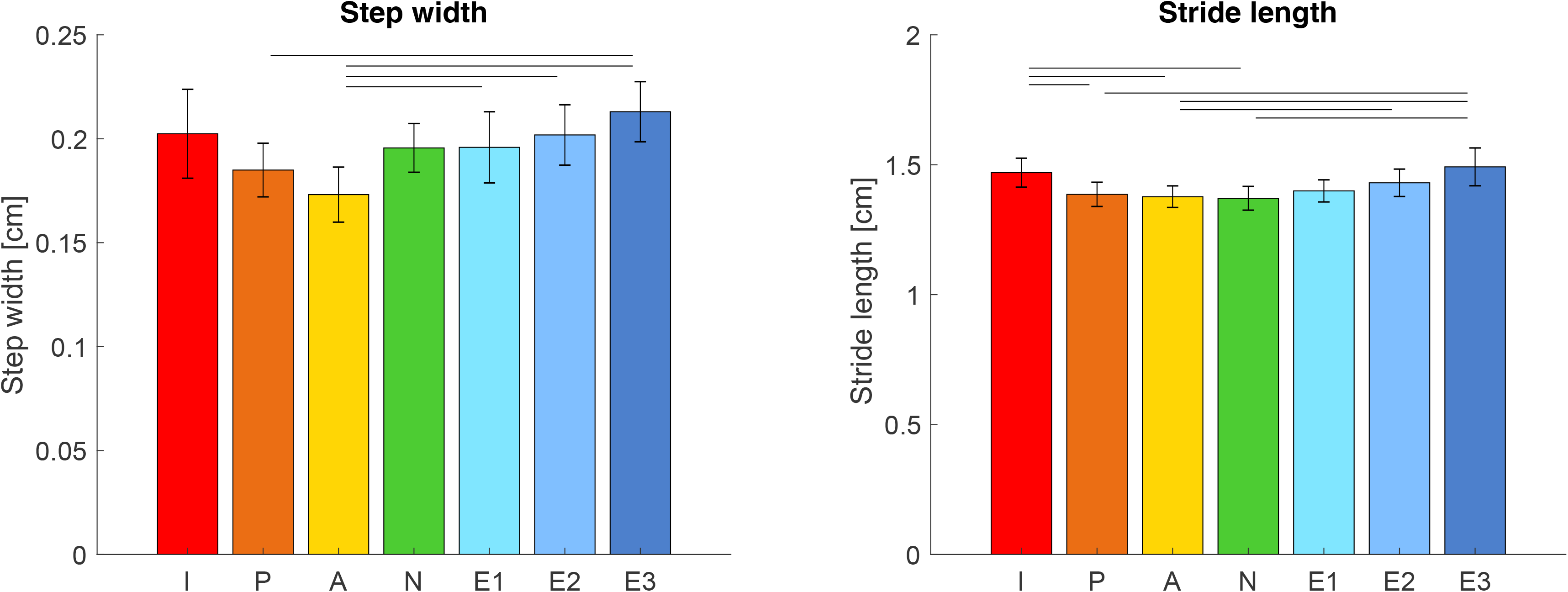
Potential confounders for the relation between VAM/GRM and Cost of Transport. (A) Shows the mean step width. (B) shows the stride frequency. (C) shows the low-frequency drift of the participant’s position. Error bars indicate the 95% Confidence Interval. Horizontal bars show significant differences between conditions (n=12, p<.05, paired t-test with bonferroni correction, and Greenhouse Geisser correction fot step width. Abbreviations: I=in-phase, P=passive (restricted), A=active (restricted), N=normal, E=extra.

Stride length was lowest in normal walking and increased as arm swing amplitude changed, resulting in a significant difference between conditions (*F*_*condition*_(6, 66)=13.85, *p*<.001).

## DISCUSSION

This study investigated the relationship between arm swing amplitude and cost of transport during walking, as well as the role of vertical angular momentum and ground reaction moment in this process. Results support the first and second hypothesis that state that when arm swing amplitude increases, VAM, and GRM decrease, albeit not for the largest arm swing amplitude. However, these changes did not always lead to an accompanying decrease in the cost of transport. Therefore, hypothesis 3 and 4 were not supported by the data.

### Influence of arm swing on VAM (hypothesis 1)

Increases in arm swing amplitude were accompanied by a decrease in whole-body VAM, in accordance with hypothesis 1, in all but one case (*extra III*). Since the arms and legs produce angular momenta opposite in sign, VAM production by the arms can compensate for VAM production at the legs. In normal walking and walking with decreased arm swing, the legs generate more momentum than the arms, leading to a net whole-body VAM unequal to zero. The extra VAM generated at the arms through the higher arm swing amplitude manages to reduce the whole-body VAM toward zero. However, it could also lead to an overcompensation and carry the VAM past zero. Further increases will then lead to an increase in VAM magnitude. In conditions where the arms overcompensated, we found a concurrent small increase in the VAM generated by the legs, which counteracted the overcompensation and kept total VAM above zero. This extra VAM from the legs may have been caused by changes in step parameters: both step width and step length increased for conditions with extra arm swing amplitude (discussed later).

Similar results for the changes in whole-body VAM were found by Collins et al. (2009a) who investigated anti-normal, held and bound arm swing (cf. *in-phase, active, and passive*) in comparison to normal walking. They found a similar pattern between conditions for the peak whole-body momentum as in-phase led to the highest VAM, normal to the lowest, and the two restricted arm swings in between. This was again due to an increase in VAM from the arms, while the contribution remained fairly constant across these conditions, similar to findings in the current study. Comparable results were also found in a study investigating walking in children with cerebral palsy. The participants showed a smaller arm swing amplitude on the affected side and higher angular momentum contributions by the legs. This was compensated by an increased arm swing on the unaffected side, so no changes in total body angular momentum were seen (Bruijn et al., 2011). To the best of our knowledge, no studies exist that investigate the influence of increased arm swing on VAM. One study (Thielemans et al., 2014) investigated the influence of adding weight to the arms, which should counter VAM by the legs in a similar way as increasing the amplitude. This study found that increasing the weight worn on the arms did not lead to a significant decrease in whole-body VAM, but this might be explained by the fact that the weight was only added to one wrist, rather than symmetrically. Thus, the participants might have compensated differently to remove asymmetries in the walking pattern or actuation thereof.

As mentioned before, we see a deviation in the general trend for extra III in the mean absolute value. Surprisingly, this increase relative to the preceding conditions is not observed in the graph of VAM is expressed as a function of the gait cycle percentage (Fig. 3A). This could be explained by the different strategies the participants used for this condition. In 5 out of 12 participants, the employed arm swing led to an overcompensation, meaning that the VAM crossed the zero and had a magnitude comparable to normal but with opposite direction. In other participants, the employed strategy actually led to an increased whole-body VAM (with the same direction as normal). Visual inspection of the walking patterns showed that some participants (#1,7 and 12) did not move their arms back all the way behind their body, rather they kept their arms in front of them thereby reducing the effectiveness of the arms in reducing the angular momentum. Participants (#4, 12) also had trouble staying in anti-phase during this condition due to different oscillation frequencies for their arms and legs, which could lead to the arms actually increasing the angular momentum in the usual direction rather than reducing it. Thus, walking with extra arm swing led to overcompensation in some participants, and increased whole-body VAM in others (see example data in Fig. 6). This led to a mean around zero when the non-absolute values over the cycle were calculated, but a higher magnitude absolute mean. For 3 out of 12 participants the whole-body VAM for *extra III* was around zero.

**Fig. 6.**
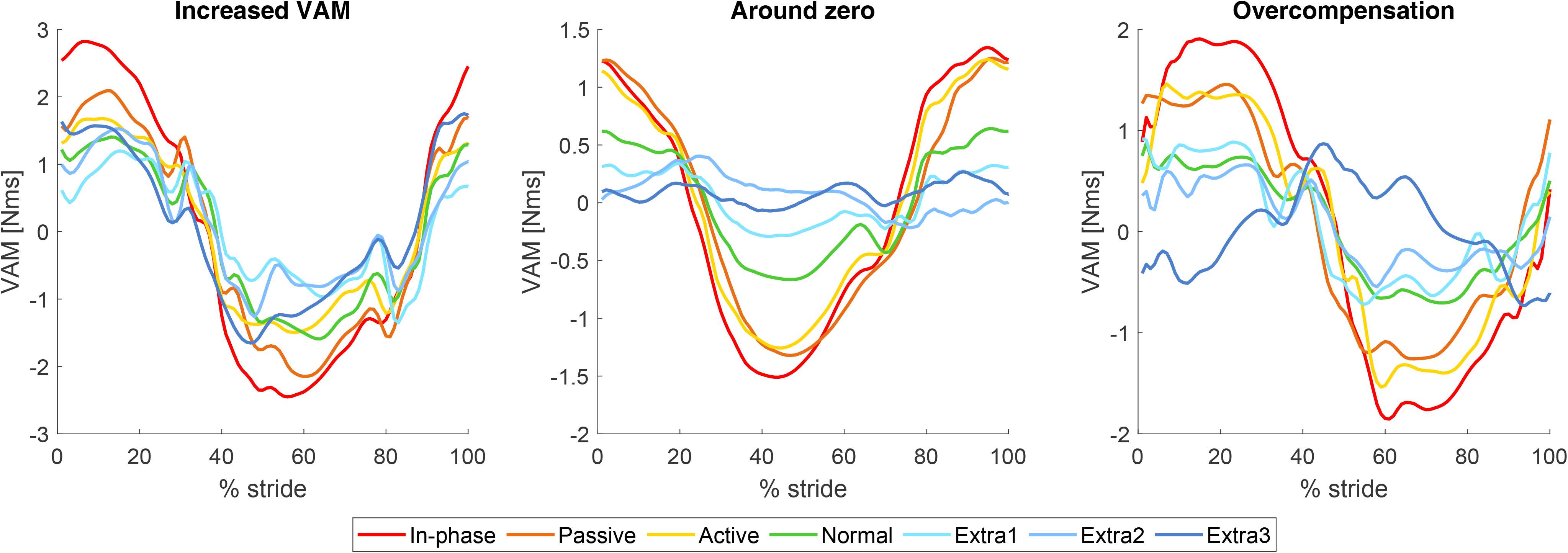
Participants used different strategies to execute the extra III condition, leading to different results for the whole-body VAM. Each of the three panels above shows one strategy to execute the extra III condition: the left panel, increased VAM, was seen in 4 participants, the results of participant #1 are shown. In the panel in the middle the participants had a whole-body VAM around zero, 3 participants used this strategy and the results of participant #10 are shown in this panel. The other 5 participants had an overcompensation due to the extra arm swing, the results of participant #5 are shown in the right panel.

### Influence of arm swing on GRM (hypothesis 2)

Similar to VAM, there was also a reduction in GRM visible for the conditions with a larger arm swing amplitude, thereby supporting the second hypothesis. This finding was not unexpected as the ground reaction moment is proportional to the time derivative of the VAM, and the VAM has a sinusoidal shape with similar periods for all conditions (N.B. the GRM is not an exact derivative in this case as they are not calculated about the same point).

The current findings are in agreement with previous studies. Collins et al. (2009a) found an increased peak vertical GRM when the hands were held or bound at the side during walking (cf. *active* and *passive* conditions), and an even further increase for in-phase walking compared to normal walking. Li et al. (2001) investigated the effect of arm fixation during walking on the ground reaction moment, and found that the GRM during walking with arm fixation (cf. passive) was significantly higher compared to normal walking in males, but not females.

### Consequences for the Cost of Transport (hypotheses 3 and 4)

The changes seen in VAM and GRM support the idea that VAM can be regulated via either arm swing or GRM. However, reducing VAM and GRM would only be favorable if these changes led to a decrease in cost of walking, as postulated in hypotheses 3 and 4. We found a pattern where the cost of transport decreased up until condition extra I (slightly more arm swing than normal). It should be noted that not all post-hoc differences were significant (see Fig. 2). When arm swing amplitude was increased beyond *extra I*, we found an increase in cost of transport, despite a reduction in VAM (from *extra I* to *extra II*) and GRM. This increase in cost of transport in the largest arm swing conditions is most likely the result of the increased cost of swinging the arms. This cost goes up as arm swing amplitude increases, due to an increasing moment arm of gravity. Furthermore, as the arm elevation goes up, the same change in arm (shoulder) angle will lead to a smaller change in horizontal amplitude (i.e. the slope of a cosine function gets smaller when approaching the peak). Taken together: the energy costs go up, while the gain goes down at larger arm elevations. These findings lead to the conclusion that hypothesis 3 and 4 should be rejected as they only hold for certain conditions. Rather, a parabolic relation between arm swing amplitude and energetic cost was found.

Notwithstanding the rejection of hypothesis 3 and 4 (for increased arm swing amplitudes), findings for reduced arm swing amplitudes agree with findings from previous studies. Collins et al. (2009a) found lowest metabolic energy for the normal walking condition, which increased 7% respectively 12% for the bound and held conditions (cf. *passive* and *active* that were +3.93% and +3.94% than normal in the current study). The cost of transport was highest in anti-normal arm swing(+26%, cf. *in-phase* which was +15.31% compared to normal in the current study) conditions. Umberger (2008) investigated the influence of walking with no arm swing on the cost of transport and found that walking with the arms folded over the chest led to a 7.7% increase in gross metabolic energy expenditure during walking. This is comparable to findings for walking without arm swing in the current study. To our knowledge, no previous studies investigated the role of increased arm swing on energetic costs so these findings cannot be compared.

### Arm swing amplitude: a trade-off?

From an evolutionary perspective, one could expect humans to walk with an optimized arm swing as there is evidence that humans optimize their walking behavior for energetic cost of transport (Holt et al., 1995; Ralston, 1958; Umberger and Martin, 2007; Zarrugh and Radcliffe, 1978). However, walking with a larger arm swing amplitude than normal did not always lead to a significant increase in energetic cost. On the contrary, the energetic cost for walking with a lightly increased arm swing (*extra I*) was even somewhat lower than for walking with normal arm swing (non-significant difference, *p*>.05). Moreover, from the viewpoint of optimizing gait stability, a larger arm swing may be beneficial (Bruijn et al., 2010; Fei Hu et al., 2012; Nakakubo et al., 2014; Punt et al., 2015). On the other hand, when faced with a larger perturbation, arm swing itself may already be detrimental (although the response of the arms to the perturbation can certainly help in recovery (Bruijn et al., 2011; Pijnappels et al., 2010)). Thus, maybe swinging the arms we do is the best trade-off between energetic cost, steady state gait stability, and maintaining the ability to respond appropriately in the face of larger perturbations.

### Study Limitations

As mentioned above, participants walked with a lower step frequency for the conditions with larger arm swing. This was not an unexpected effect since it seems logical that if the arm swing amplitude increases, so will the step length to keep the velocity of arm swing in a preferable range. With a constant speed imposed by the treadmill this means that step frequency will go down. The change in step length might also be a reaction to the overcompensation of the VAM through the arms, in order to counteract it. In either case, the changes in these step parameters can potentially influence current findings. It has been shown that individuals tend to walk with a preferred speed-frequency relation and that deviation from this optimal relation can lead to an increase in cost of transport (Bertram and Ruina, 2001). Therefore, the cost of transport for the *extra arm* swing amplitudes and *in-phase* are potentially higher due to participants walking with a non-optimal step frequency. Step length also appears to influence VAM, with smaller steps leading to a lower whole-body VAM (Thielemans et al., 2014), when walking with normal arm swing amplitude. Beside the changes in step frequency and step length, the step width also changed during the different arm swing conditions, becoming larger in the non-normal arm swing conditions. These changes can have an independent influence on the cost of transport. Donelan et al. (2001) found a 45% higher energetic cost of transport when people walk with a wider step width compared to their preferred step width, and an 8% higher energetic cost for walking with a smaller step width. Thus, the higher cost of transport found in the normal arm swing conditions might be in part due to the wider step width that people walk with, in these conditions.

All participants walked with a constant average speed of 1.25 m/s. The speed of walking has been shown to have an effect on the vertical angular momentum in both human experiments (Thielemans et al., 2014) as well as in modeling studies (Collins et al., 2009a). Therefore, the effect of increasing arm swing could be different for other walking speeds.

## CONCLUSION

This study explored the relation between arm swing amplitude, vertical angular momentum, ground reaction moment and cost of transport by having participants walk with different styles and amplitudes of arm swing. Our findings support the hypotheses that VAM and GRM decrease with increasing arm swing amplitude (resp. hypotheses 1 and 2). The decrease in total VAM is the result of the increase VAM contribution of the arms, that can now compensate for a larger part of the VAM generated by the legs. In some cases, this led to an overcompensation.

Cost of transport was optimal around normal and slightly increased arm swing amplitudes. The hypothesis that the reduced VAM and GRM lead to a decreased cost of transport was confirmed up until this optimal point. Increasing arm swing beyond that led to an increased cost of transport, most likely due to the disproportional increase in cost of swinging the arms.

In conclusion, increasing arm swing amplitude leads to a reduction in vertical angular moment and ground reaction moments. However, this is not always useful in terms of cost of transport, which is congruent with the evolutionary concept of metabolically optimized walking. It might, however, provide useful if one wants to decrease the ground reaction moment, for instance to alleviate the legs in lower extremity disorders. Normal or slightly increased arm swing amplitude appears to be optimal in young and healthy individuals. This natural arm swing might be the best trade-off between energetic cost, steady state gait stability, and the ability to respond to larger gait perturbations.

## ACKNOWLEDGEMENTS

SMB was funded by a VIDI grant (016.Vidi.178.014) from the Dutch Organization for Scientific Research (NWO).

## LIST OF SYMBOLS AND ABBREVIATIONS

*a*: Distance (m)
ap: anteroposterior
COM: Center of Mass
COP: Center of Pressure
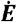: Energy consumption in (W) or (W kg^−1^)
*g*: Gravitational constant
GRF: Ground Reaction Force (N)
GRM: Ground Reaction Moment (Nm)
*I*: Inertia tensor
*l*: Leg length (m)
*L*: Angular Momentum (Nms)
*m*: mass (kg)
ml: Mediolateral
*M*_*z*_: Total vertical moment around the origin of the force platform
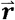: Position Vector (m)
RER: Respiratory Exchange Ratio
RM-ANOVA: Repeated Measures Analysis of Variance
*s*: Number of segments
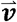: Velocity Vector
VAM: Vertical Angular Momentum (Nms)
VO_2_: Oxygen Uptake (ml s^−1^)
*ω*: Angular velocity (rad s^−1^)

## REFERENCES

Bertram, J. E. A. and Ruina, A. (2001). Multiple walking speed-frequency relations are predicted by constrained optimization. J. Theor. Biol. 209, 445–453.

Bruijn, S. M., Meijer, O. G., van Dieën, J. H., Kingma, I. and Lamoth, C. J. C. (2008). Coordination of leg swing, thorax rotations, and pelvis rotations during gait: The organisation of total body angular momentum. Gait Posture 27, 455–462.

Bruijn, S. M., Meijer, O. G., Beek, P. J. and van Dieën, J. H. (2010). The effects of arm swing on human gait stability. J. Exp. Biol. 213,.

Bruijn, S. M., Meyns, P., Jonkers, I., Kaat, D. and Duysens, J. (2011). Control of angular momentum during walking in children with cerebral palsy. Res. Dev. Disabil. 32, 2860–2866.

Collins, S. H., Adamczyk, P. G. and Kuo, A. D. (2009a). Dynamic arm swinging in human walking. Proc. Biol. Sci. 276, 3679–88.

Collins, S. H., Adamczyk, P. G. and Kuo, A. D. (2009b). Dynamic arm swinging in human walking. Proc. Biol. Sci. 276, 3679–88.

Donelan, J. M., Kram, R. and Kuo, A. D. (2001). Mechanical and metabolic determinants of the preferred step width in human walking. Proc. R. Soc. B Biol. Sci. 268, 1985–1992.

Elftman, H. (1939). The Function of the Arms in Walking. Hum. Biol. 11, 529–535.

Fei Hu, Dong-Yun Gu, Jin-Ling Chen, Yu Wu, Bing-Chen An and Ke-Rong Dai (2012). Contribution of arm swing to dynamic stability based on the nonlinear time series analysis method. In 2012 Annual International Conference of the IEEE Engineering in Medicine and Biology Society, pp. 4831–4834. IEEE.

Fernandez-Ballesteros, M. L., Buchthal, F., Rosenfalck, P., Ballesteros, M. L. F., Buchthal, F. and Rosenfalck, P. (1965). The Pattern of Muscular Activity During the Arm Swing of Natural Walking. Acta Physiol. Scand. 63, 296–310.

Garby, L. and Astrup, A. (1987). The relationship between the respiratory quotient and the energy equivalent of oxygen during simultaneous glucose and lipid oxidation and lipogenesis. Acta Physiol. Scand. 129, 443–444.

Gerdy, P. N. (1829). Memoires sur le mecanisme de la marche de l’homme. J. Physiol. expérimentale Pathol. 9, 1–28.

Goudriaan, M., Jonkers, I., Van Dieen, J. H. and Bruijn, S. M. (2014). Arm swing in human walking: What is their drive?

Herr, H. and Popovic, M. (2008). Angular momentum in human walking. J. Exp. Biol. 211, 467–481.

Hinrichs, R. N. (1990). Whole body movement: coordination of arms and legs in walking and running. Mult. Muscle Syst. springer New York 694–705.

Hof, A. L. (2007). The equations of motion for a standing human reveal three mechanisms for balance. J. Biomech. 40, 451–457.

Holt, K. G., Jeng, S. F., Ratcliffe, R. and Hamill, J. (1995). Energetic Cost and Stability during Human Walking at the Preferred Stride Frequency. J. Mot. Behav. 27, 164–178.

Jackson, K. M., Joseph, J. and Wyard, S. J. (1978). A mathematical model of arm swing during human locomotion. J. Biomech. 11, 277–289.

Kubo, M., Wagenaar, R. C., Saltzman, E. and Holt, K. G. (2004). Biomechanical mechanism for transitions in phase and frequency of arm and leg swing during walking. Biol. Cybern. 91, 91–8.

Kuhtz-Buschbeck, J. P. and Jing, B. (2012). Activity of upper limb muscles during human walking. J. Electromyogr. Kinesiol. 22, 199–206.

Li, Y., Wang, W., Crompton, R. H. and Gunther, M. M. (2001). Free vertical moments and transverse forces in human walking and their role in relation to arm-swing. J. Exp. Biol. 204, 47–58.

Marigold, D. S., Bethune, A. J. and Patla, A. E. (2002). Role of the Unperturbed Limb and Arms in the Reactive Recovery Response to an Unexpected Slip During Locomotion. J. Neurophysiol. 89, 1727–1737.

Meyns, P., Bruijn, S. M. and Duysens, J. (2013). The how and why of arm swing during human walking. Gait Posture 38, 555–562.

Morton, D. J. and Fuller, D. D. (1952). Human locomotion and body form□:a study of gravity and man. Baltimore: The Williams & Wilkins Company.

Murray, M. P., Sepic, S. B. and Barnard, E. J. (1967). Patterns of sagittal rotation of the upper limbs in walking. Phys. Ther. 47, 272–84.

Nakakubo, S., Doi, T., Sawa, R., Misu, S., Tsutsumimoto, K. and Ono, R. (2014). Does arm swing emphasized deliberately increase the trunk stability during walking in the elderly adults? Gait Posture 40, 516–520.

Ortega, J. D., Fehlman, L. A. and Farley, C. T. (2008). Effects of aging and arm swing on the metabolic cost of stability in human walking. J. Biomech. 41, 3303–8.

Park, J. (2008). Synthesis of natural arm swing motion in human bipedal walking. J. Biomech. 41, 1417–1426.

Pijnappels, M., Kingma, I., Wezenberg, D., Reurink, G. and van Dieën, J. H. (2010). Armed against falls: The contribution of arm movements to balance recovery after tripping. Exp. Brain Res. 201, 689–699.

Pontzer, H., Holloway, J. H., Raichlen, D. A. and Lieberman, D. E. (2009). Control and function of arm swing in human walking and running. J. Exp. Biol. 212, 523–534.

Punt, M., Bruijn, S. M., Wittink, H. and van Dieën, J. H. (2015). Effect of arm swing strategy on local dynamic stability of human gait. Gait Posture 41, 504–509.

Ralston, H. J. (1958). Energy-speed relation and optimal speed during level walking. Int. Zeitschrift für Angew. Physiol. Einschl. Arbeitsphysiologie 17, 277–283.

Thielemans, V., Meyns, P. and Bruijn, S. M. (2014). Is angular momentum in the horizontal plane during gait a controlled variable? Hum. Mov. Sci. 34, 205–216.

Umberger, B. R. (2008). Effects of suppressing arm swing on kinematics, kinetics, and energetics of human walking. J. Biomech. 41, 2575–2580.

Umberger, B. R. and Martin, P. E. (2007). Mechanical power and efficiency of level walking with different stride rates. J. Exp. Biol. 210, 3255–3265.

Weber, W. and Weber, E. (1836). Mechanik der menschlichen Gehwerkzeuge. Dietrichsche Buchhandlung.

Witte, H., Preuschoft, H. and Recknagel, S. (1991). Human body proportions explained on the basis of biomechanical principles. Z. Morphol. Anthropol. 78, 407–23.

Zarrugh, M. Y. and Radcliffe, C. W. (1978). Predicting metabolic cost of level walking. Eur. J. Appl. Physiol. Occup. Physiol. 38, 215–223.

Zehr, E. P. and Duysens, J. (2004). Regulation of arm and leg movement during human locomotion. Neuroscientist 10, 347–61.

